# Recognizability bias in citizen science photographs

**DOI:** 10.1101/2022.06.25.497604

**Authors:** Wouter Koch, Laurens Hogeweg, Erlend B. Nilsen, Robert B. O’Hara, Anders G. Finstad

## Abstract

Citizen science initiatives and automated collection methods increasingly depend on image recognition in order to provide the amounts of observational data research and management needs. Training recognition models, meanwhile, also requires large amounts of data from these sources, creating a feedback loop between the methods and the tools. Species that are harder to recognize, both for humans and machine learning algorithms, are likely to be underreported, and thus be less prevalent in the training data. As a result, the feedback loop may hamper training mostly for species that already pose the greatest challenge. In this study, we trained recognition models for various taxa, and found evidence for a *“recognizability bias”*, where species that models struggle with are also generally underreported. This has implications for the kind of performance one can expect from future models that are trained with more data, including such challenging species. We consider identification methods that rely on more than photographs alone to be important in improving future identification tools.

## Introduction

There is an ever growing need for large amounts of biodiversity observation data. With an increasing awareness of the multiple crises biodiversity faces [1–3], substantial amounts of such data are essential if humanity is to monitor trends and address these issues [4–6]. Occurrence data are typically subject to spatial, temporal and taxonomic bias [7, 8], and traditional manual methods of data collection are insufficient to gather the data volume needed, or address these biases. Alternative data collection methods, ranging from citizen science (nonprofessional volunteers reporting observations [9]) to camera-traps automating insect monitoring [10, 11] are being deployed to gather large amounts of data. With the increased output from such initiatives, manual management and quality control become infeasible. Automated image recognition tools for species identification are increasingly used to facilitate this [12–15]. Training image recognition models, however, also requires large amounts of pictures [16]. This creates a mutual reliance between large scale image data collection and image recognition models [17].

Visual identification of species is a complex task, and taxa vary in their recognizability; while some species are unmistakable, many others are very challenging or even outright impossible to identify, regardless of picture quality [18]. As models are trained using training data reported and identified by humans, species with low recognizability among humans will be underreported and be underrepresented in the training data. This affects recognition models, as these are then being trained with data biased towards higher recognizability, consisting mostly of pictures of species that are easier to recognize. If this is the case, training models will be hampered not only by the lower recognizability of particularly challenging species, but also by their higher absence from the training data.

To evaluate the existence of this possible bias and its consequences, we evaluated how data availability, picture quality, biological traits and data collection differs across species within 3 orders of birds, and how these differences relate to recognition model performance. All data came from a large Norwegian citizen science project, where recognition tools are not a part of the reporting or validation process. Birds are the most well-represented orders per species, allowing for the most detailed analysis. We also trained models for 9 other orders of plants, animals and fungi, to test for a general correlation between data availability and model performance, and to evaluate what this means for future recognition models.

We find evidence for a *“recognizability bias”*, where species that are more readily identified by humans and recognition models alike are more prevalent in the available image data. This pattern is present across multiple taxa, and does not appear to relate to a difference in picture quality, biological traits, or data collection metrics other than recognizability.

## Methods

We trained image recognition models using convolutional neural networks on pictures retrieved from the Norwegian citizen science platform Species Observation Service [19] for 12 orders: Agaricales, Anseriformes, Asparagales, Asterales, Charadriiformes, Coleoptera, Diptera, Lecanorales, Lepidoptera, Odonata, Passeriformes, and Polyporales [20]. A separate model was trained for each order, using 200 documented observations per species for training and validation, and a minimum of 20 for the test set. See Koch *et al*. [21] for details. From these models and various external datasets, several relevant metrics were collected (table 1).

**Table 1:**
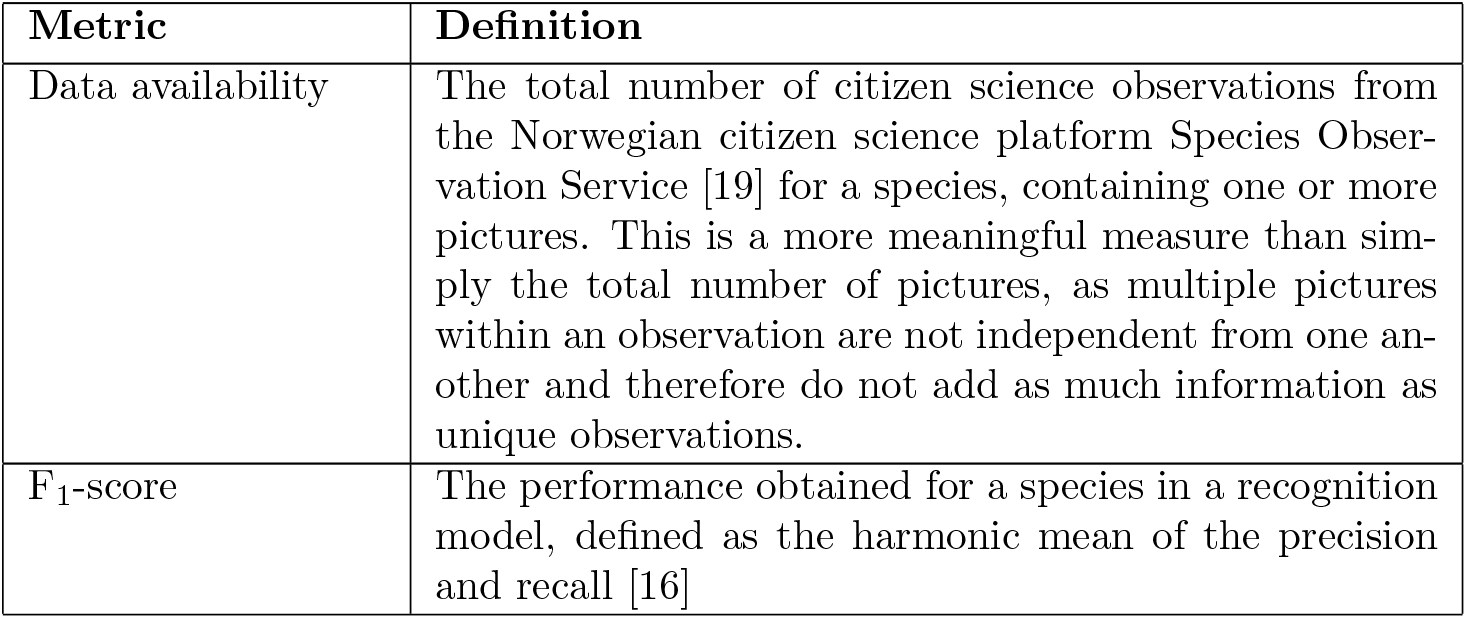
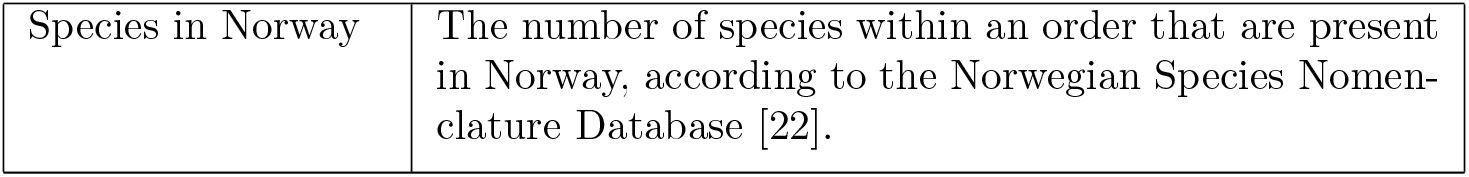
Metrics collected for species within all orders

More detailed analyses were done on the included bird orders; waterfowl (Anseriformes), shorebirds (Charadriiformes), and passerines (Passeriformes), as bird orders have the highest proportion of species in Norway represented in the dataset, and ample standardized available data on a range of biological traits allowing for a deeper analysis. For these analyses, a number of additional metrics were collected for the included bird species (table 2).

**Table 2:** Metrics collected for species within the bird orders

| Metric | Definition |
| --- | --- |
| Picture quality | Using Label Studio v1.4 [23], $\geq 50$ pictures per species were annotated by drawing rectangles approximately equal in surface area to the visible part of each individual bird. From this, we took the percentage of the picture occupied by the largest depiction of an individual of the target species, minus the percentage of the picture occupied by all individuals of other bird species. Per species, the median log value was used as a proxy for picture quality. |
| Urbanness | The proportion of 100 documented observations from the Species Observation Service with a location within a cell tagged as “urban” in the ESA CCI landcover dataset [24]. |
| Hand-wing index | Wing length minus wing width, a measure positively correlated with flight efficiency and dispersal ability of a species. Retrieved from the Global-HWI dataset [25]. |
| Body mass | The average log-transformed body mass of a species, retrieved from the Global-HWI dataset [25]. |
| Habitat openness | A three-step scale of the openness of the habitat of a species, retrieved from the Global-HWI dataset [25]. |
| Documentation rate | The proportion, per species, of observations in the Species Observation Service that have one or more pictures. |
| Picture density | The average number of pictures per observation from the Species Observation Service, from those with at least one picture. |
| Observation rate | The number of observations in the Species Observation Service dataset per observation in the TOV-e bird monitoring scheme [26] |

LASSO multiple regression models were trained using Scikit-learn [27] to evaluate the effect of the biological traits, picture quality measurement, and data collection process from table 2 on the F_1_-scores for birds. All LASSO models have the order as a factor. The full model for biological traits is given by

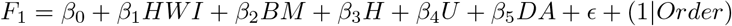

where *HWI* is the hand-wing index, *BM* is the body mass, *H* is the habitat openness, *U* is the urbanness, and *DA* is the log data availability. The full model for picture quality is given by

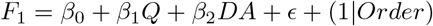

where *Q* is the picture quality, and *DA* is the log data availability. The full model for data collection parameters is given by

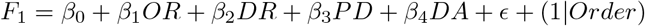

where *OR* is the observation rate, *DR* is the documentation rate, *PD* is the picture density, and *DA* is the log data availability.

## Results

There is a strong positive linear correlation between log data availability and the F_1_-score for bird species (figure 1). Note that data availability does not affect training, as all models were trained and evaluated using 220 documented observations per species, regardless of the total availability. A positive linear correlation was also evident in 7 of the 9 other orders (figure 2), in particular Asterales and Odonata. The beetles (Coleoptera) and lichens (Lecanorales) exhibited no apparent correlation, with an R^2^ of 0.06 and 0.12, and P-values of 0.27 and 0.18, respectively.

**Figure 1:**
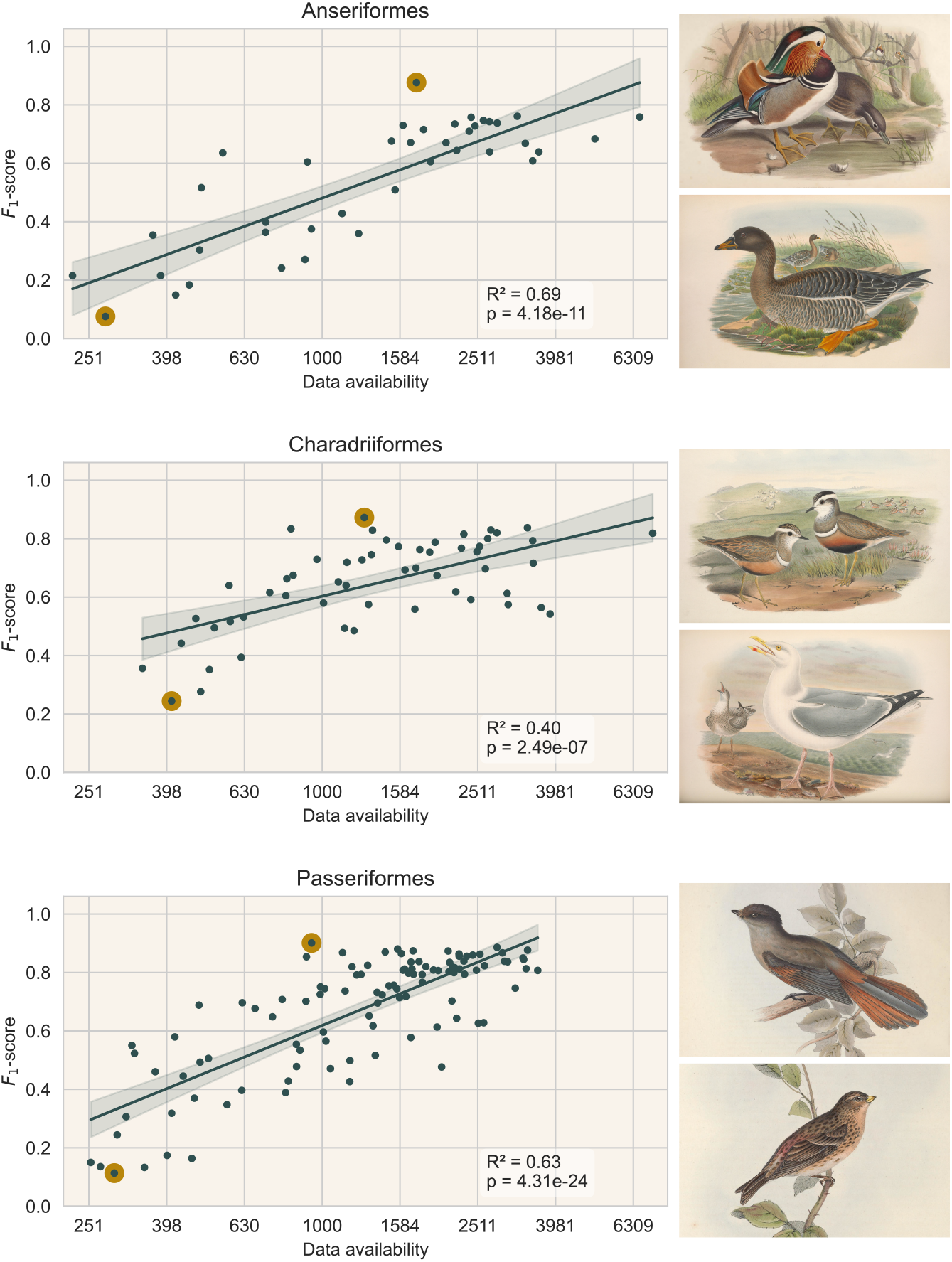
Effect of the total data availability per species on their F_1_-scores, in models trained with 200 documented observations, for three bird orders. The topand bottom-performing species per order (highlighted dots) are depicted, see table S1. Regressions are Ordinary Least Squares with 95% confidence intervals.

**Figure 2:**
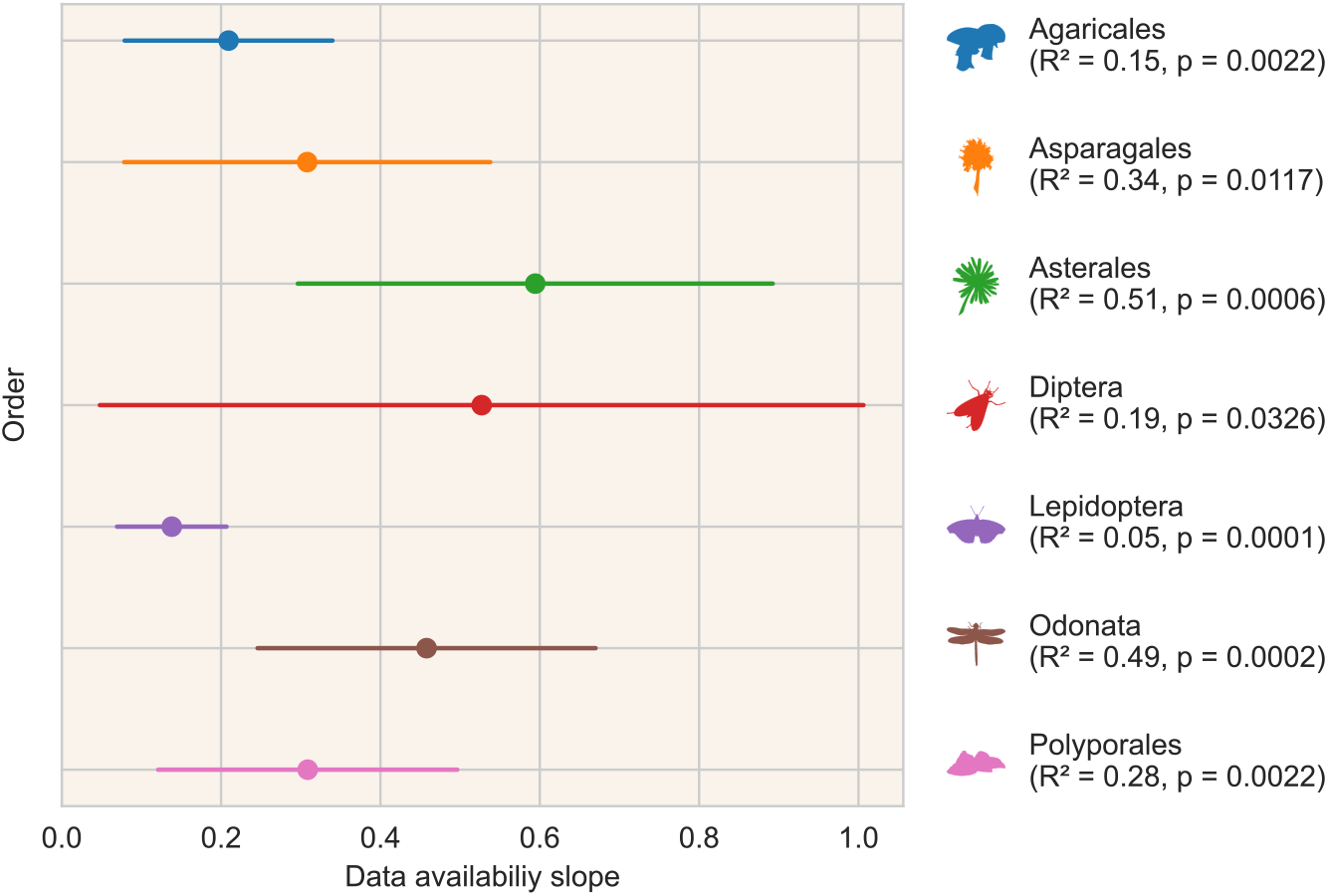
The slopes of the correlations between total data availability per species and their F_1_-scores, in models trained with 200 documented observations, for non-bird orders with a correlation p*<*0.05. Regressions are Ordinary Least Squares, lines indicate the 95% confidence intervals.

In each bird order, there is a linear relationship between species’ picture density and documentation rate (R^2^≥0.52, p≤1.51×10^*−*7^, see table S2). We also find a negative linear correlation between picture density and F_1_-scores (R^2^≥0.23, p≤2.1×10^−4^, see table S2), and some negative linear correlation between documentation rate and F_1_-scores (R^2^≥0.11, p≤4.64×10^−3^, see table S2). For passerines, there is a negative linear relationship between habitat openness and picture quality (R^2^ = 0.26, p = 3.53×10^−8^, see table S2). Waterfowl and shorebirds could not be evaluated as they only occur in open habitats.

LASSO models trained on biological traits, collection process parameters, and picture quality, all having and log data availability as an additional parameter and order as a factor, had R^2^ values of 0.60, 0.57 and 0.63, respectively.

With that, none of the full model performances were substantial improvements from a LASSO model with log data availability as its only parameter (R^2^ = 0.57).

## Discussion

We find a conspicuous pattern where recognition models attain higher performances for species that are reported with pictures more frequently. It is probable that the recognizability of the species influences both their likelihood of being reported with pictures, as well as recognition model performances. The citizen science project used as a data source here does not include any recognition tools in its reporting or validation process, allowing a distinction between human and algorithm recognition biases. Unmistakable species can be recognized and reported by more citizen scientists, resulting in greater data availability for such species. A recognition model, dealing with the same information as human observers, is also proportionally more likely to reliably recognize these species.

This is supported by a qualitative comparison between species with the highest and lowest recognition model performances, where easy to recognize, characteristic species are reported more often than hard to recognize species (e.g. nondescript species or species similar to other related species) (see figure 1 and table S1). Further support comes from the fact that most of the correlation is explained by the data availability for a species, rather than the documentation rate or the picture density. Thus, there is more data available mainly when a species is recognized and reported more, rather than it being disproportionately more likely to be reported with pictures, or with many pictures when reported with pictures.

An alternative explanation to recognizability for increased model performance might be a difference in the kind of pictures, but we find no evidence for this. Species traits, habitat use, and image quality could affect recognition model performance if pictures of more photographed birds are taken more up close, with higher zoom, or were cropped more. We found no evidence, however, for a link between model performance and either picture quality or biological traits in birds. For the passerines, where habitat openness varies among species, we do find that picture quality decreases for species associated with more open habitats. It makes intuitive sense that birds in open habitats are photographed from a greater distance than their forest dwelling counterparts, which will be hidden from view unless in close proximity. While this intercorrelation supports the validity of the picture quality metric, neither habitat nor picture quality affect recognition model performance. We conclude that differences in model performance are caused by the recognizability of the species, rather than by how, or how large species are generally depicted.

Since multiple pictures connected to a single observation are not truly independent, training data are generated based on the number of documented observations, rather than the total number of pictures. One might expect that species with a higher picture density will perform better, as observations with more pictures can provide some additional information in the training process. We find a reverse effect however, where performance for such species is substantially lower. A likely explanation is that species with high picture densities are rarities in Norway (e.g. the top 3 species being Caspian gull, Blyth’s reed warbler, and Pine bunting). Species with the lowest picture density, meanwhile, are typical common, well-known species such as corvids and titmice. Rarities are reported not because they are easy to find or identify by casual observers, but due to their popularity among avid birdwatchers, who are likely to document their observations. A strong correlation between picture density and documentation rate supports this; rarities are more often reported with pictures, and in such cases relatively often with several pictures.

While we investigated the bird orders in detail, the link between data availability and model performance is present in other orders too (figure 2). Some orders are notoriously difficult to identify to species level, e.g. flies (Diptera) and beetles (Coleoptera), but our models for these perform surprisingly well. The list of species with sufficient observations with pictures for inclusion in the experiment reveals that only relatively easy to recognize species, often with distinct colorations (e.g. ladybugs for beetles) are represented in this subset.

More generally, the requirement that species must have at least 220 citizen science observations with pictures generates a non-random subset of species, and it differs greatly per order how selective this criterion is. Bird species are most frequently reported; 48% of the species present in Norway [22] within the bird orders examined here meet the selection criterion. One of the other orders for which the pattern was found, the dragonflies and damselflies (Odonata), have only 52 species in Norway, of which 44% met the criteria for inclusion. This is in stark contrast to the beetles (1% inclusion), and lichens (2% inclusion), where no clear correlation is found. It is reasonable to assume that for these taxa, the experiment only considers the most recognizable species. If observations were thousandfold, more challenging species could be included, giving a broader range in performances and possibly a similar positive correlation between model performance and data availability.

The consequence of the recognizability bias found here is that as more data is collected, ultimately providing the numbers of pictures needed to train models also on less reported, harder to recognize species, current performance of recognition models cannot be extrapolated to these expanded models. In other words, data that are lacking now are in part lacking because such species are harder to recognize. When such data is added in the future, the performance increase will not be as great as in the past. Besides citizen science, even methods that have no inherent reporting bias, such as automated insect camera traps and trail cameras, can still be subject to recognizability bias. There too, species that are less readily identified will result in more unidentifiable pictures, providing relatively less training data.

Image recognition tools play an important role in maintaining the quality of the large amounts of biodiversity data science and management require. There are limits to what can be identified from a picture however, and identification tools are needed that rely on more than just pixel information. Models that take into account season, location, sound, etc. can be especially beneficial for difficult species. Still, there is no substitute for the taxonomic knowledge of experts. Preserving this knowledge, and making it available in the form of identification keys is vital. These can be powerful tools to more reliably identify challenging species, in tandem with automatic identification.

## Acknowledgements

We are grateful to Rune Sørås, Ingeborg H. Bringslid, and Rienk W. Fokkema for their help in annotating pictures.

## Data accessibility

All code is available through Zenodo at https://doi.org/10.5281/zenodo. 6734696. Bird illustrations in figure 1 are works in the Public Domain made by John Gould (1804-1881), obtained through the Biodiversity Heritage Library [28–30]

## Authors’ contributions

WK: conception, experimental design, code, analysis, writing. LH: code, text revision. EBN: conception, text revision. RBOH: analysis, text revision. AGF: conception, analysis, text revision.

