## Supplementary information for "Recognizability bias in citizen science photographs"

| **Order** | **Species** | **$F_1$-score** |
|---|---|---|
| Passeriformes | *Perisoreus infaustus* | 0.901 |
| Passeriformes | *Cinclus cinclus* | 0.886 |
| Passeriformes | *Periparus ater* | 0.88 |
| Passeriformes | *Bombycilla garrulus* | 0.876 |
| Anseriformes | *Aix galericulata* | 0.876 |
| Passeriformes | *Certhia familiaris* | 0.874 |
| Passeriformes | *Aegithalos caudatus* | 0.873 |
| Charadriiformes | *Charadrius morinellus* | 0.872 |
| Passeriformes | *Regulus regulus* | 0.87 |
| Passeriformes | *Lophophanes cristatus* | 0.868 |
| Passeriformes | *Emberiza citrinella* | 0.867 |
| Passeriformes | *Garrulus glandarius* | 0.865 |
| Passeriformes | *Pyrrhula pyrrhula* | 0.863 |
| Passeriformes | *Pinicola enucleator* | 0.863 |
| Passeriformes | *Cyanistes caeruleus* | 0.86 |
| Passeriformes | *Sitta europaea* | 0.856 |
| Passeriformes | *Turdus merula* | 0.855 |
| Passeriformes | *Phylloscopus sibilatrix* | 0.854 |
| Passeriformes | *Coccothraustes coccothraustes* | 0.85 |
| Passeriformes | *Carduelis carduelis* | 0.845 |
| Passeriformes | *Motacilla cinerea* | 0.839 |
| Passeriformes | *Erithacus rubecula* | 0.838 |
| Passeriformes | *Parus major* | 0.837 |
| Charadriiformes | *Haematopus ostralegus* | 0.837 |
| Passeriformes | *Motacilla alba* | 0.837 |
| Passeriformes | *Prunella modularis* | 0.836 |
| Passeriformes | *Lanius collurio* | 0.835 |
| Charadriiformes | *Phalaropus lobatus* | 0.834 |
| Charadriiformes | *Calidris maritima* | 0.83 |
| Charadriiformes | *Arenaria interpres* | 0.829 |
| Passeriformes | *Phylloscopus inornatus* | 0.824 |
| Passeriformes | *Saxicola rubicola* | 0.823 |
| Charadriiformes | *Charadrius hiaticula* | 0.82 |
| Passeriformes | *Sylvia atricapilla* | 0.82 |
| Passeriformes | *Turdus philomelos* | 0.819 |
| Charadriiformes | *Gallinago gallinago* | 0.819 |
| Passeriformes | *Luscinia svecica* | 0.818 |
| Passeriformes | *Plectrophenax nivalis* | 0.816 |
| Charadriiformes | *Tringa totanus* | 0.815 |
| Passeriformes | *Lanius excubitor* | 0.813 |

| | | |
|---|---|---|
| Passeriformes | *Turdus viscivorus* | 0.812 |
| Passeriformes | *Fringilla coelebs* | 0.812 |
| Passeriformes | *Nucifraga caryocatactes* | 0.812 |
| Passeriformes | *Passer montanus* | 0.808 |
| Passeriformes | *Turdus iliacus* | 0.808 |
| Passeriformes | *Emberiza schoeniclus* | 0.808 |
| Passeriformes | *Oenanthe oenanthe* | 0.807 |
| Passeriformes | *Saxicola rubetra* | 0.807 |
| Passeriformes | *Chloris chloris* | 0.802 |
| Charadriiformes | *Calidris alpina* | 0.8 |
| Passeriformes | *Anthus petrosus* | 0.8 |
| Passeriformes | *Motacilla flava* | 0.798 |
| Charadriiformes | *Cepphus grylle* | 0.795 |
| Passeriformes | *Fringilla montifringilla* | 0.794 |
| Passeriformes | *Troglodytes troglodytes* | 0.794 |
| Charadriiformes | *Vanellus vanellus* | 0.793 |
| Passeriformes | *Carpodacus erythrinus* | 0.793 |
| Passeriformes | *Turdus torquatus* | 0.793 |
| Passeriformes | *Eremophila alpestris* | 0.792 |
| Charadriiformes | *Actitis hypoleucos* | 0.788 |
| Charadriiformes | *Pluvialis apricaria* | 0.773 |
| Charadriiformes | *Tringa glareola* | 0.773 |
| Charadriiformes | *Limosa lapponica* | 0.767 |
| Passeriformes | *Ficedula hypoleuca* | 0.766 |
| Charadriiformes | *Charadrius dubius* | 0.762 |
| Anseriformes | *Cygnus olor* | 0.761 |
| Anseriformes | *Mergellus albellus* | 0.758 |
| Anseriformes | *Clangula hyemalis* | 0.757 |
| Passeriformes | *Muscicapa striata* | 0.756 |
| Charadriiformes | *Calidris pugnax* | 0.755 |
| Passeriformes | *Phoenicurus ochruros* | 0.754 |
| Charadriiformes | *Tringa nebularia* | 0.754 |
| Passeriformes | *Acrocephalus schoenobaenus* | 0.751 |
| Anseriformes | *Branta leucopsis* | 0.747 |
| Passeriformes | *Sturnus vulgaris* | 0.746 |
| Charadriiformes | *Limosa limosa* | 0.745 |
| Passeriformes | *Calcarius lapponicus* | 0.745 |
| Passeriformes | *Phoenicurus phoenicurus* | 0.744 |
| Anseriformes | *Mergus merganser* | 0.742 |
| Anseriformes | *Bucephala clangula* | 0.738 |
| Passeriformes | *Curruca communis* | 0.737 |
| Anseriformes | *Tadorna tadorna* | 0.734 |
| Passeriformes | *Poecile montanus* | 0.732 |

| | | |
|---|---|---|
| Anseriformes | *Melanitta fusca* | 0.73 |
| Charadriiformes | *Tringa erythropus* | 0.729 |
| Anseriformes | *Anas acuta* | 0.728 |
| Charadriiformes | *Calidris minuta* | 0.727 |
| Passeriformes | *Poecile palustris* | 0.725 |
| Passeriformes | *Passer domesticus* | 0.723 |
| Charadriiformes | *Calidris temminckii* | 0.719 |
| Passeriformes | *Hirundo rustica* | 0.718 |
| Charadriiformes | *Numenius arquata* | 0.716 |
| Anseriformes | *Mareca penelope* | 0.715 |
| Passeriformes | *Corvus frugilegus* | 0.714 |
| Anseriformes | *Mergus serrator* | 0.71 |
| Passeriformes | *Sylvia curruca* | 0.708 |
| Passeriformes | *Turdus pilaris* | 0.703 |
| Passeriformes | *Pica pica* | 0.702 |
| Charadriiformes | *Uria aalge* | 0.699 |
| Charadriiformes | *Chroicocephalus ridibundus* | 0.697 |
| Passeriformes | *Hippolais icterina* | 0.696 |
| Passeriformes | *Loxia leucoptera* | 0.696 |
| Charadriiformes | *Calidris canutus* | 0.693 |
| Passeriformes | *Emberiza pusilla* | 0.688 |
| Anseriformes | *Cygnus cygnus* | 0.683 |
| Passeriformes | *Panurus biarmicus* | 0.677 |
| Anseriformes | *Somateria spectabilis* | 0.676 |
| Charadriiformes | *Alca torda* | 0.675 |
| Charadriiformes | *Rissa tridactyla* | 0.674 |
| Anseriformes | *Branta canadensis* | 0.671 |
| Anseriformes | *Aythya fuligula* | 0.67 |
| Anseriformes | *Anas platyrhynchos* | 0.668 |
| Charadriiformes | *Alle alle* | 0.663 |
| Charadriiformes | *Calidris alba* | 0.652 |
| Passeriformes | *Alauda arvensis* | 0.651 |
| Passeriformes | *Corvus monedula* | 0.648 |
| Anseriformes | *Anas crecca* | 0.644 |
| Passeriformes | *Phylloscopus collybita* | 0.643 |
| Charadriiformes | *Pluvialis squatarola* | 0.64 |
| Charadriiformes | *Fratercula arctica* | 0.64 |
| Anseriformes | *Somateria mollissima* | 0.639 |
| Anseriformes | *Anser anser* | 0.639 |
| Anseriformes | *Anser indicus* | 0.635 |
| Passeriformes | *Acanthis flammea* | 0.628 |
| Passeriformes | *Anthus pratensis* | 0.627 |
| Charadriiformes | *Sterna hirundo* | 0.618 |

| | | |
|---|---|---|
| Passeriformes | *Corvus cornix* | 0.618 |
| Charadriiformes | *Stercorarius parasiticus* | 0.616 |
| Passeriformes | *Phylloscopus trochilus* | 0.613 |
| Charadriiformes | *Larus hyperboreus* | 0.613 |
| Anseriformes | *Anser brachyrhynchus* | 0.608 |
| Anseriformes | *Aythya marila* | 0.605 |
| Charadriiformes | *Calidris ferruginea* | 0.605 |
| Anseriformes | *Aythya ferina* | 0.605 |
| Passeriformes | *Corvus corax* | 0.596 |
| Charadriiformes | *Larus fuscus* | 0.592 |
| Charadriiformes | *Tringa ochropus* | 0.58 |
| Passeriformes | *Locustella naevia* | 0.579 |
| Passeriformes | *Carduelis spinus* | 0.577 |
| Charadriiformes | *Numenius phaeopus* | 0.575 |
| Charadriiformes | *Larus canus* | 0.574 |
| Passeriformes | *Carduelis flavirostris* | 0.565 |
| Charadriiformes | *Larus glaucoides* | 0.564 |
| Charadriiformes | *Larus marinus* | 0.559 |
| Passeriformes | *Anthus trivialis* | 0.554 |
| Passeriformes | *Regulus ignicapilla* | 0.55 |
| Charadriiformes | *Larus argentatus* | 0.542 |
| Passeriformes | *Riparia riparia* | 0.534 |
| Charadriiformes | *Scolopax rusticola* | 0.532 |
| Charadriiformes | *Phalaropus fulicarius* | 0.526 |
| Passeriformes | *Sylvia nisoria* | 0.523 |
| Anseriformes | *Polysticta stelleri* | 0.517 |
| Passeriformes | *Acanthis hornemanni* | 0.516 |
| Charadriiformes | *Stercorarius skua* | 0.516 |
| Anseriformes | *Melanitta nigra* | 0.509 |
| Passeriformes | *Sylvia borin* | 0.506 |
| Passeriformes | *Loxia pytyopsittacus* | 0.498 |
| Charadriiformes | *Calidris falcinellus* | 0.495 |
| Charadriiformes | *Sterna paradisaea* | 0.493 |
| Passeriformes | *Lullula arborea* | 0.493 |
| Charadriiformes | *Hydrocoloeus minutus* | 0.485 |
| Passeriformes | *Pastor roseus* | 0.478 |
| Passeriformes | *Loxia curvirostra* | 0.477 |
| Passeriformes | *Acanthis cabaret* | 0.471 |
| Passeriformes | *Turdus atrogularis* | 0.46 |
| Passeriformes | *Ficedula parva* | 0.445 |
| Charadriiformes | *Stercorarius longicaudus* | 0.442 |
| Passeriformes | *Corvus corone* | 0.428 |
| Anseriformes | *Anas clypeata* | 0.428 |

| | | |
|---|---|---|
| Passeriformes | *Carduelis cannabina* | 0.426 |
| Anseriformes | *Anser albifrons* | 0.399 |
| Passeriformes | *Acrocephalus dumetorum* | 0.397 |
| Charadriiformes | *Calidris melanotos* | 0.394 |
| Passeriformes | *Acrocephalus palustris* | 0.389 |
| Anseriformes | *Anas strepera* | 0.375 |
| Passeriformes | *Delichon urbicum* | 0.37 |
| Anseriformes | *Mareca strepera* | 0.364 |
| Anseriformes | *Anser fabalis* | 0.359 |
| Charadriiformes | *Thalasseus sandvicensis* | 0.356 |
| Anseriformes | *Branta bernicla* | 0.354 |
| Charadriiformes | *Larus melanocephalus* | 0.352 |
| Passeriformes | *Acrocephalus scirpaceus* | 0.347 |
| Passeriformes | *Motacilla citreola* | 0.318 |
| Passeriformes | *Luscinia luscinia* | 0.307 |
| Anseriformes | *Aythya collaris* | 0.303 |
| Charadriiformes | *Lymnocryptes minimus* | 0.276 |
| Anseriformes | *Spatula clypeata* | 0.271 |
| Passeriformes | *Emberiza leucocephalos* | 0.244 |
| Charadriiformes | *Larus cachinnans* | 0.244 |
| Anseriformes | *Anas querquedula* | 0.241 |
| Anseriformes | *Anas carolinensis* | 0.215 |
| Anseriformes | *Tadorna ferruginea* | 0.215 |
| Anseriformes | *Cygnus columbianus* | 0.184 |
| Passeriformes | *Anthus richardi* | 0.174 |
| Passeriformes | *Spinus spinus* | 0.164 |
| Passeriformes | *Anthus hodgsoni* | 0.15 |
| Anseriformes | *Spatula querquedula* | 0.149 |
| Passeriformes | *Anthus cervinus* | 0.136 |
| Passeriformes | *Linaria cannabina* | 0.133 |
| Passeriformes | *Linaria flavirostris* | 0.113 |
| Anseriformes | *Anser serrirostris* | 0.076 |

Table S1: Metrics collected for species within the bird orders

| **Dependent variable** | **Parameters** | **Slope** | **Intercept** | **$R^2$** | **P-value** |
|---|---|---|---|---|---|
| Agaricales $F_1$-score | Data availability (log) | 0.21 | 0.27 | 0.15 | $2.15\times10^{-3}$ |
| Anseriformes documentation rate | Picture density | 0.38 | -0.47 | 0.52 | $1.51\times10^{-7}$ |

| | | | | | |
|---|---|---|---|---|---|
| Anseriformes $F_1$-score | Data availability (log) | 0.48 | -0.97 | 0.69 | $4.18\times10^{-11}$ |
| Anseriformes $F_1$-score | Documentation rate | -1.52 | 0.63 | 0.19 | $4.64\times10^{-3}$ |
| Anseriformes $F_1$-score | Picture density | -1.02 | 1.95 | 0.31 | $2.10\times10^{-4}$ |
| Asparagales $F_1$-score | Data availability (log) | 0.31 | 0.01 | 0.34 | 0.0117 |
| Asterales $F_1$-score | Data availability (log) | 0.59 | -0.71 | 0.51 | $5.92\times10^{-4}$ |
| Charadriiformes documentation rate | Picture density | 0.34 | -0.43 | 0.76 | $5.51\times10^{-18}$ |
| Charadriiformes $F_1$-score | Data availability (log) | 0.32 | -0.34 | 0.4 | $2.49\times10^{-7}$ |
| Charadriiformes $F_1$-score | Documentation rate | -0.9 | 0.7 | 0.28 | $3.45\times10^{-5}$ |
| Charadriiformes $F_1$-score | Picture density | -0.42 | 1.25 | 0.39 | $3.52\times10^{-7}$ |
| Coleoptera $F_1$-score | Data availability (log) | 0.13 | 0.57 | 0.06 | 0.273 |
| Diptera $F_1$-score | Data availability (log) | 0.53 | -0.63 | 0.19 | 0.0326 |
| Lecanorales $F_1$-score | Data availability (log) | 0.2 | 0.31 | 0.12 | 0.118 |
| Lepidoptera $F_1$-score | Data availability (log) | 0.14 | 0.49 | 0.05 | $9.50\times10^{-5}$ |
| Odonata $F_1$-score | Data availability (log) | 0.46 | -0.53 | 0.49 | $2.02\times10^{-4}$ |
| Passeriformes documentation rate | Picture density | 0.3 | -0.36 | 0.55 | $1.59\times10^{-19}$ |
| Passeriformes $F_1$-score | Data availability (log) | 0.54 | -1 | 0.63 | $4.31\times10^{-24}$ |
| Passeriformes $F_1$-score | Documentation rate | -0.85 | 0.71 | 0.11 | $5.19\times10^{-4}$ |
| Passeriformes $F_1$-score | Picture density | -0.5 | 1.37 | 0.23 | $1.68\times10^{-7}$ |
| Passeriformes picture quality | Habitat openness | -0.12 | 5.93 | 0.26 | $5.53\times10^{-8}$ |
| Polyporales $F_1$-score | Data availability (log) | 0.31 | -0.11 | 0.28 | $2.18\times10^{-3}$ |

Table S2: Metrics collected for species within the bird orders
